## Supplemental figures for "Cryo-EM Structures of CTP Synthase Filaments Reveal Mechanism of pH-Sensitive Assembly During Budding Yeast Starvation"

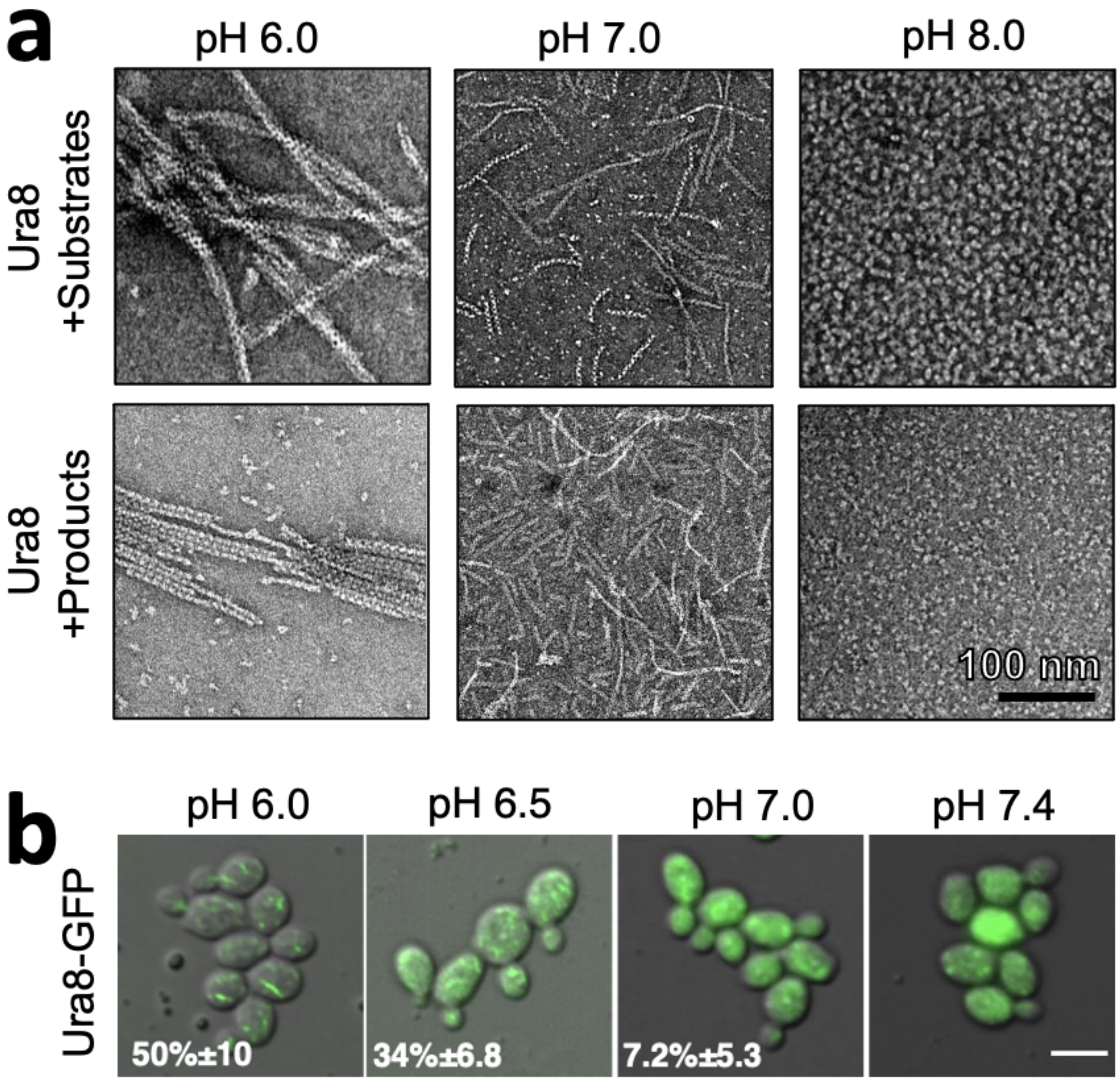


**Fig2. Supplemental 1. Ura8 assembly is driven by pH and substrates or products. (a)** Negative stain EM of purified recombinant Ura8 assembled with substrates (2mM UTP/2mM ATP) or product (2mM CTP). **(b)** Yeast expressing GFP-tagged Ura8 with membrane permeabilization using 2mM DNP and supplemented with 2% glucose. Scale bar is 5 microns.

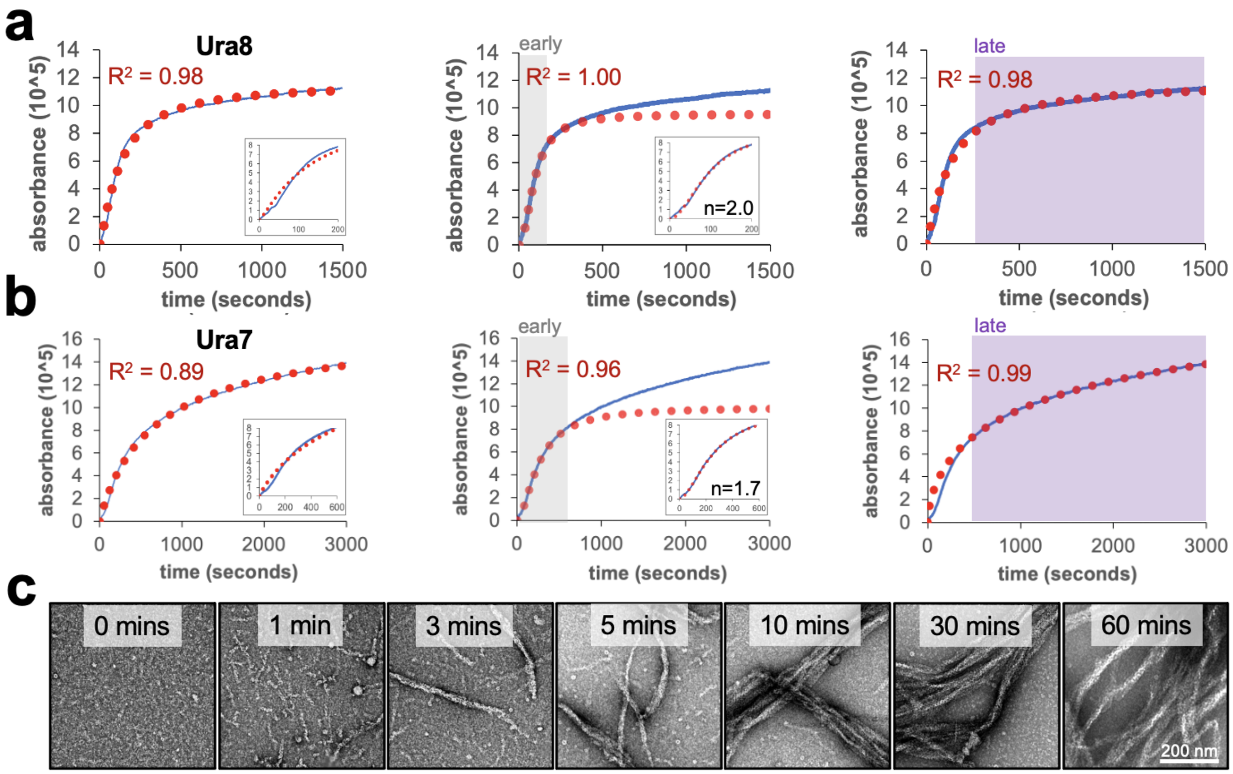


**Fig2. Supplemental 2. Right angle light scattering assembly kinetics for CTPS filaments. (a)** Right angle light scattering (blue line) for product-driven Ura8 assembly at pH 6.0. A single four-parameter logistic regression (red dotted line) fit to the full data (left), early (middle; 0-300sec) or late (right; 300-1500sec) time-points. Insets show zoom in of early time points. $R^2$ is the square of the Pearson product moment correlation coefficient, representing the quality of the fit. **(b)** Same as panel A but for Ura7. **(c)** Negative stain EM of Ura7 assembly from panel B.

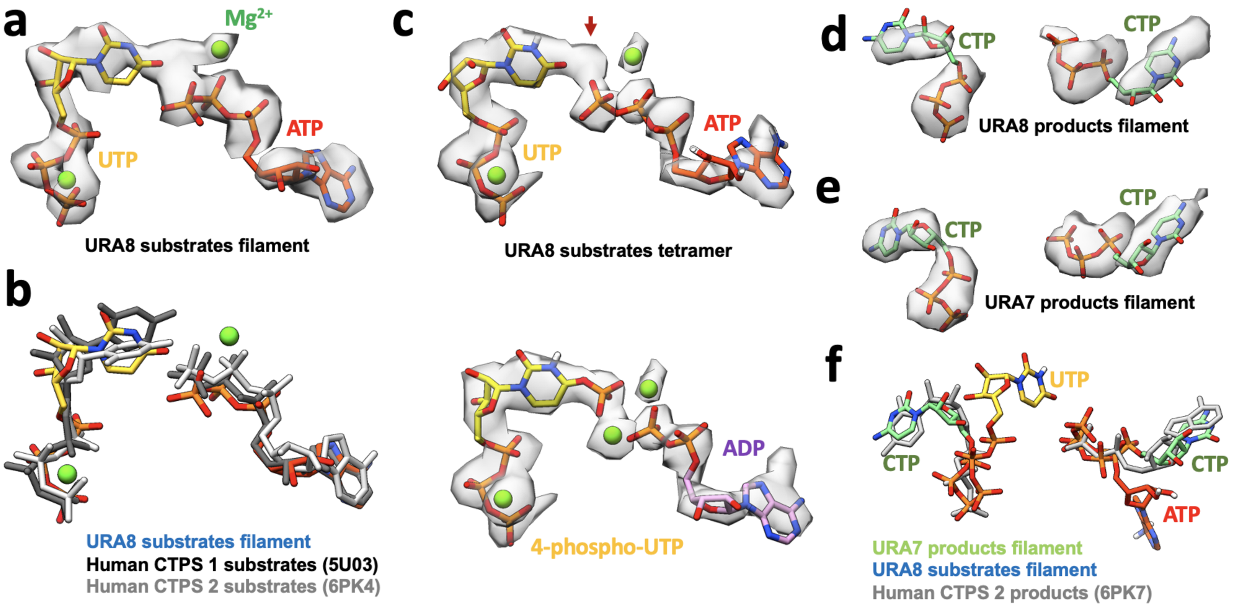


**Fig3. Supplemental 1. Nucleotides in yeast CTPS. (a)** Ura8 bound to ATP/UTP in filament with corresponding map density. **(b)** comparison of model from panel A to published human CTPS 1 and 2 bound to substrates. **(c)** Ura8 bound to ATP/UTP in tetramer with corresponding map density. Arrow shows extra unknown density. Bottom panel shows the same density modelled with hydrolyzed ATP, yielding a 4-phospho-UTP intermediate in the active site. **(d-e)** CTP-bound URA8 and URA7 in filaments with corresponding density. **(f)** Overlay of product-bound Ura7 from panel E with published product-bound human CTPS2. Substrate-bound Ura8 filament ATP and UTP are displayed to show relative positioning of products and substrates in the active site.

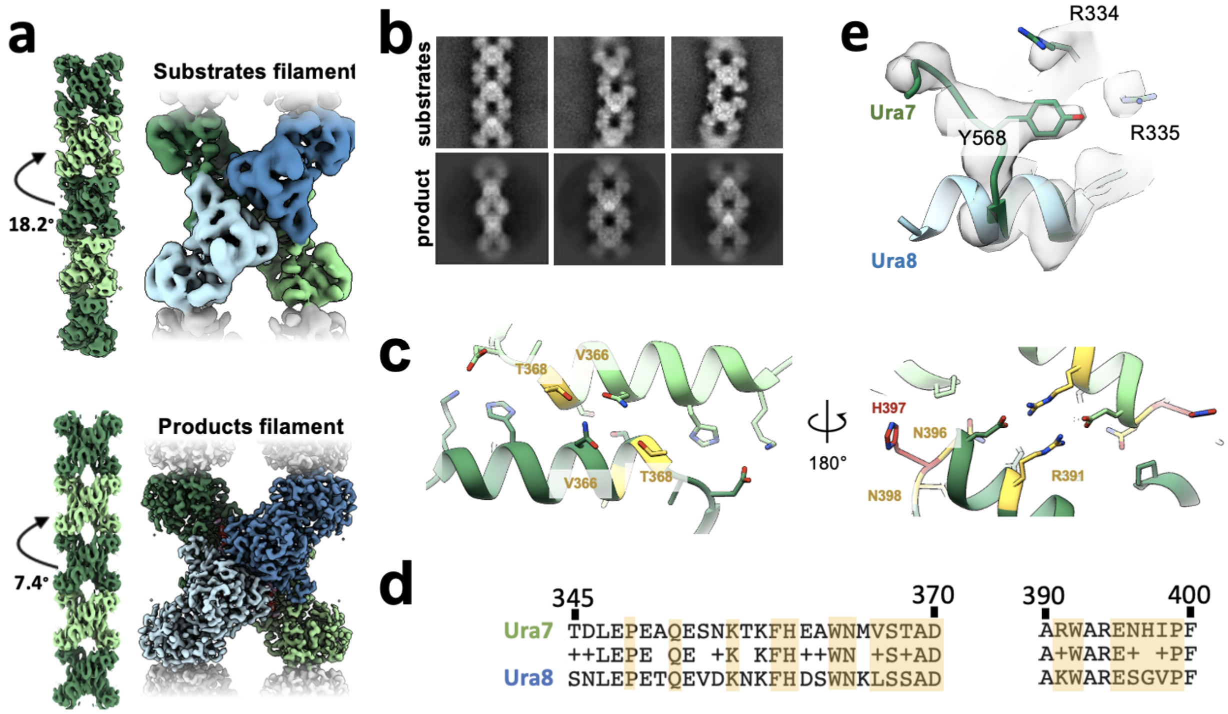


**Fig3. Supplemental 2. Ura7 filament architecture and assembly interface.** **(a)** cryo-EM reconstructions of Ura7 in substrate and product bound states. Left are maps generated from imposing helical symmetry parameters on a reconstruction of a single protomer. **(b)** Selected 2D class averages of substrate- or product-bound Ura7. **(c)** Ura7 filament interface. Residues painted similar (yellow) and different (red) from Ura8. In addition, Ura7 assembles with the W363/W392 hydrophobic core shown in figure 3. **(d)** Sequence alignment of filament assembly interface for Ura7 (top) and Ura8 (bottom). Residues participating in interface assembly are highlighted in yellow.

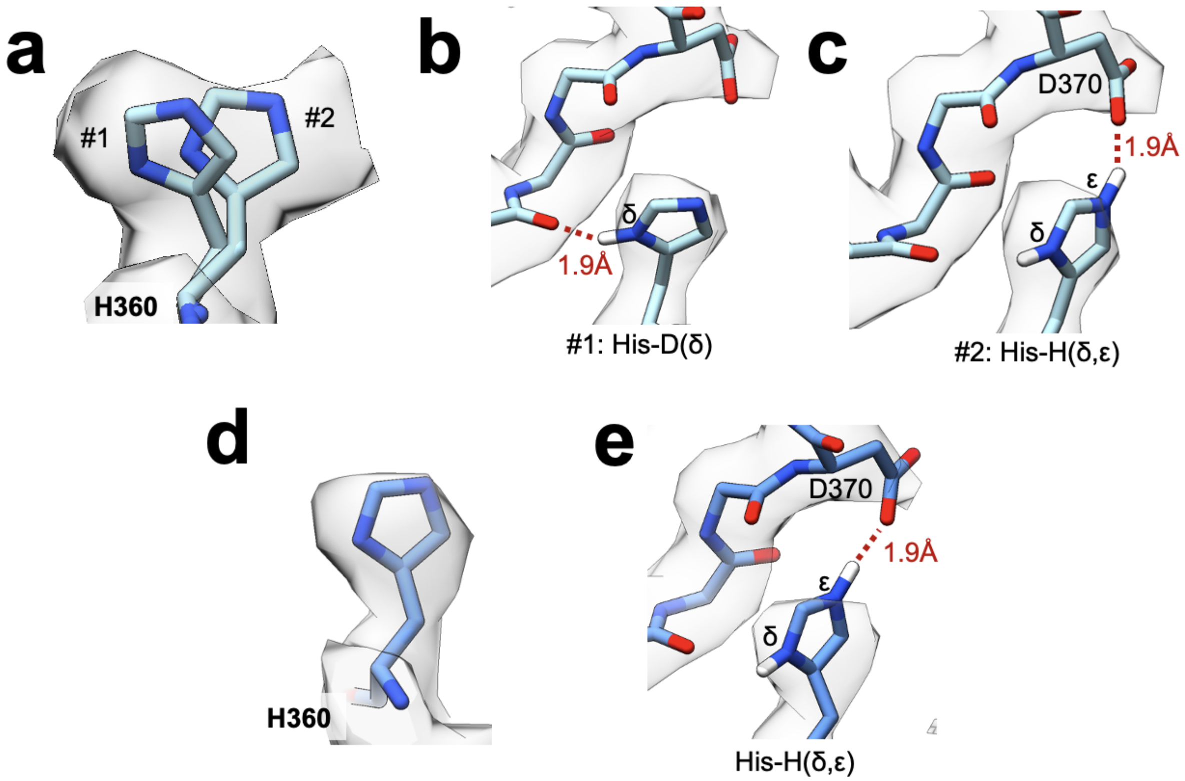


**Fig 3 supp 3. Protonation State at the Filament Assembly interface.** **(a)** Substrate-bound Ura8 filament map density for H360 with two possible rotamers modelled. Note that this view is rotated slightly relative to panels b-c. **(b)** Single-protonation state (His-D) of H360 hydrogen of rotamer #1 from panel A bonding with a backbone carbonyl across the interface. **(c)** Double-protonation state (His-H) of rotamer #2 from panel A driven by low pH leads to an ionic interaction of epsilon hydrogen with D370. **(d)** Similar to panel A, for His360 from Substrate-bound Ura8 bundle. **(e)** Double-protonation state of H360 making an ionic interaction with D370 in substrate-bound Ura8 bundle.

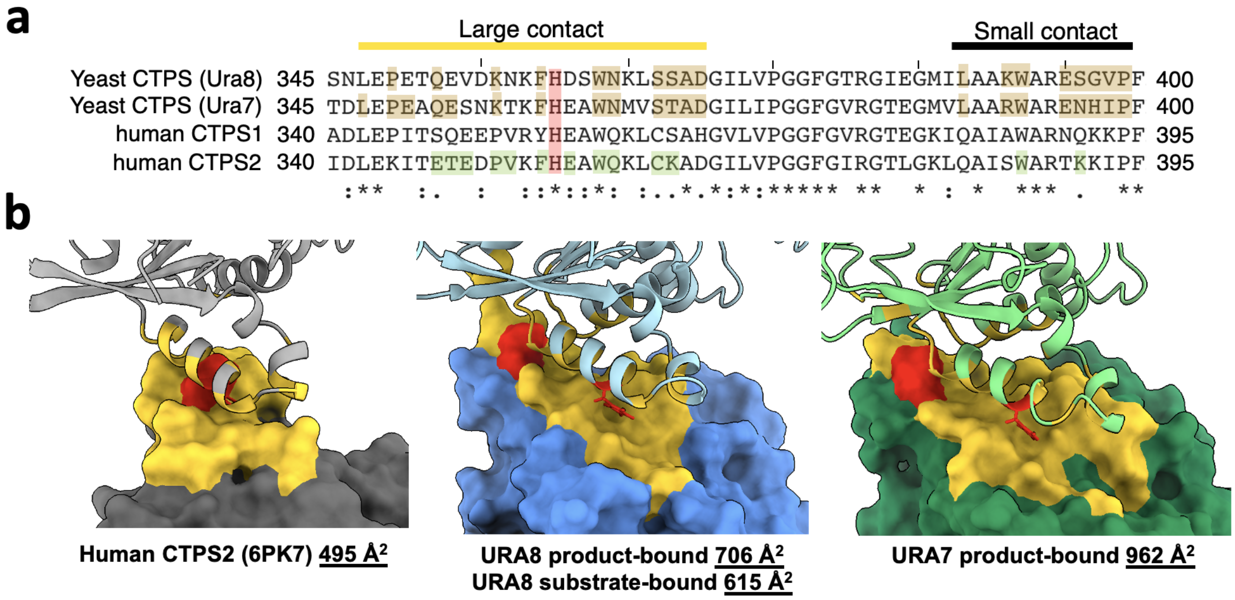


**Fig3. Supplemental 4. Comparison of buried surface area for yeast and human CTPS. (a)** Sequence alignment of filament assembly interface for human and yeast isoforms. Residues highlighted in brown and green correspond to filament assembly contacts for yeast and human ctps, respectively. Conserved histidine highlighted in red. **(b)** Calculated buried surface area for one monomer of human or yeast CTPS with interfacing residues painted yellow. H360 (H355 in humans) is colored in red.

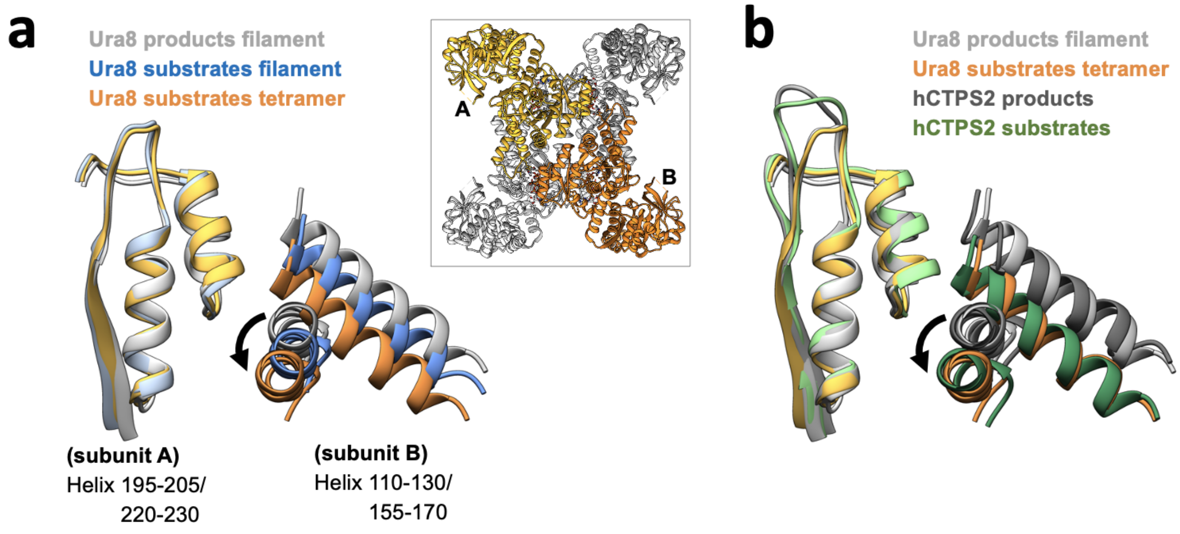


**Fig3. Supplemental 5. Amido-ligase domain movement at the tetramerization interface (a)** helices at the tetramerization interface for a transverse pair of monomers (A and B in inset). Models aligned on subunit A helices (left) to show relative movement of subunit B. **(b)** models of yeast CTPS (Ura8) helices from panel A shown relative to published human CTPS2 (6PK4 substrates; 6PK7 products).

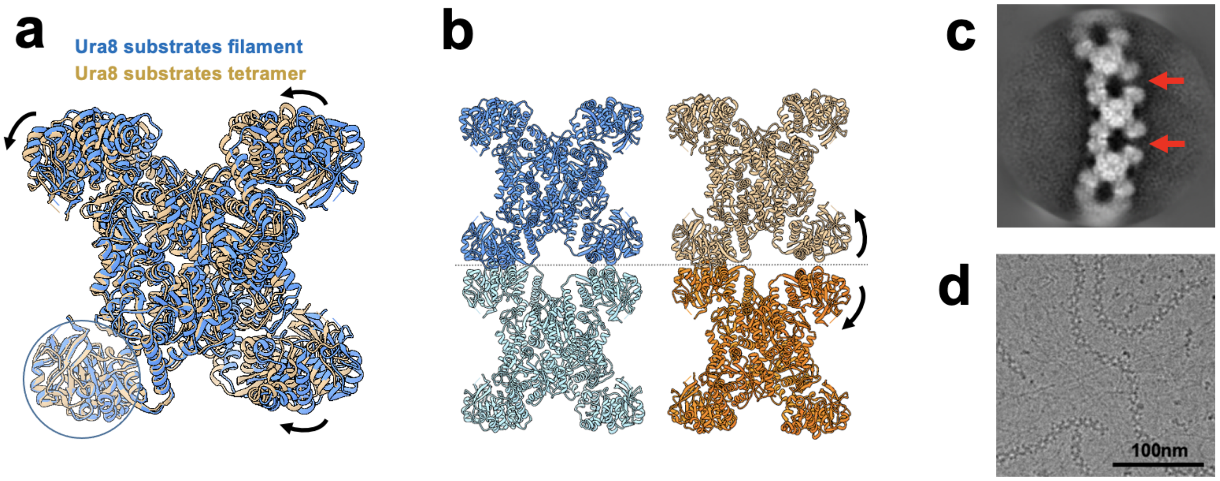


**Fig3. Supplemental 6. CTPS filaments are incompatible with the active state** **(a)** Tetramer models for substrate-bound Ura8 in a filament (low pH) or unassembled (high pH). Models aligned on the glutaminase domain for one monomer (circled) and the resulting movement is shown with arrows. **(b)** substrate-bound Ura8 filament (blue, left) used for alignment of two substrate-bound tetramers (right) on the contacting glutaminase domains as done in panel A. When tetramers are aligned on the left glutaminase domain, the opposing (right) interface splays open, disrupting filament architecture. **(c)** Representative 2D class average of substrate-bound Ura7 showing breaking of filaments described in panel B. **(d)** Example cryo-EM micrograph of substrate-bound Ura7 at low pH.

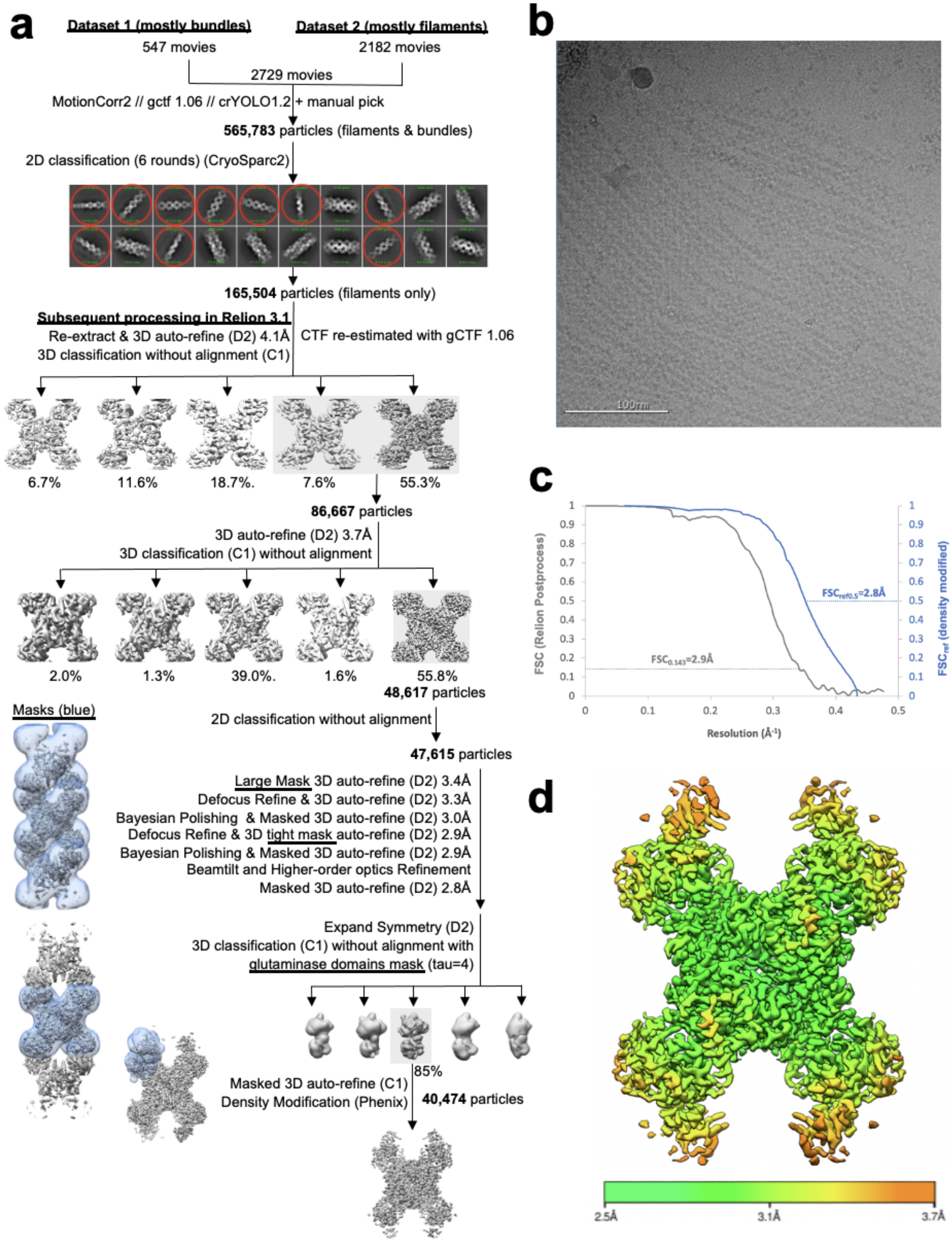


**Fig3. Supplemental 7. Image Processing of Substrate-bound Ura8 Filaments.** (a) Flowchart overview of the data processing strategy. (b) example micrograph from the dataset. (c) FSC curve from Relion postprocess and Phenix density modification. (d) local resolution estimation.

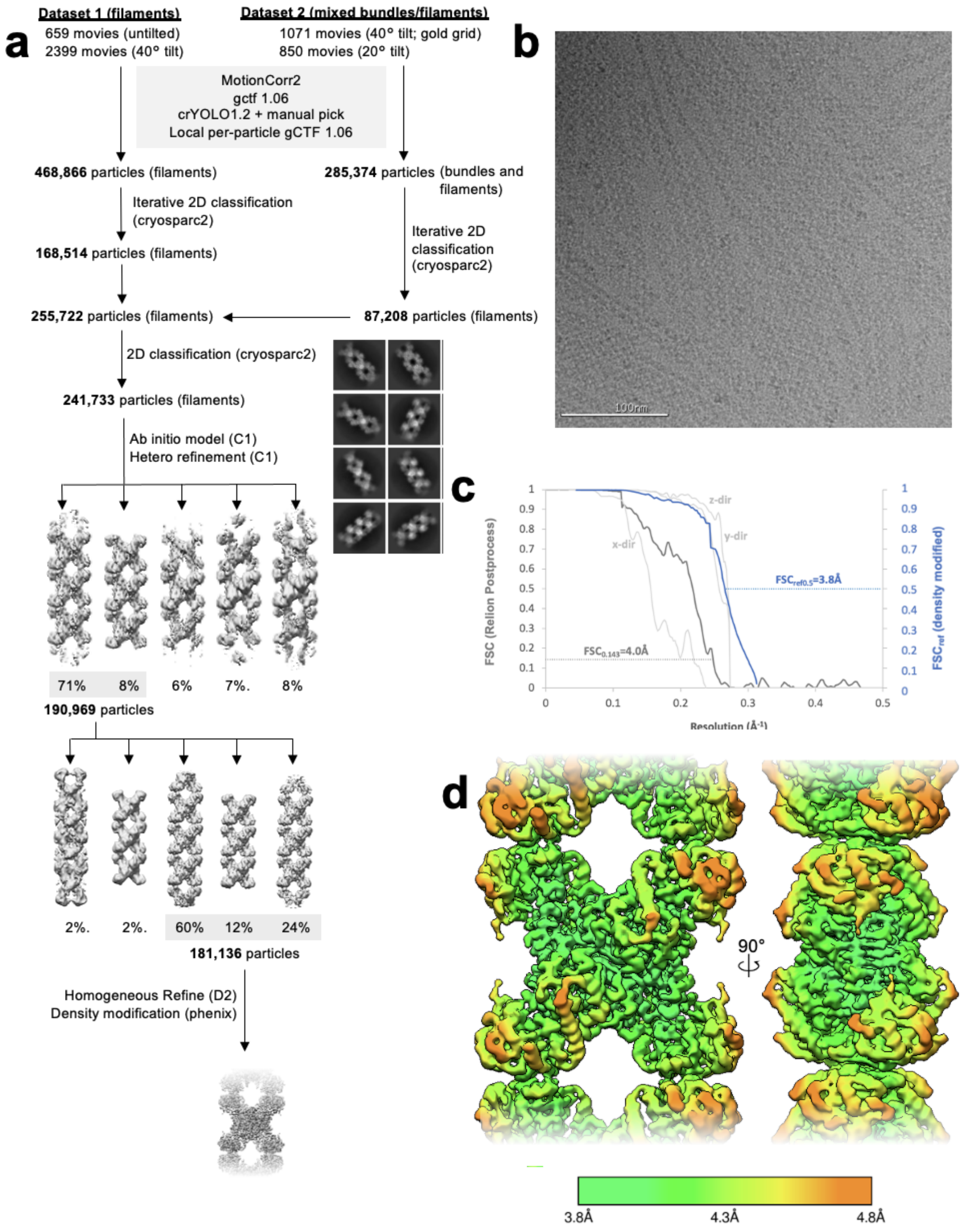


**Fig3. Supplemental 8. Image Processing of Product-bound Ura8 Filaments.** (a) Flowchart overview of the data processing strategy. (b) example micrograph from the dataset. (c) FSC curve from Relion postprocess and Phenix density modification. Light grey is 3D FSC for X/Y/Z directions. (d) local resolution estimation.

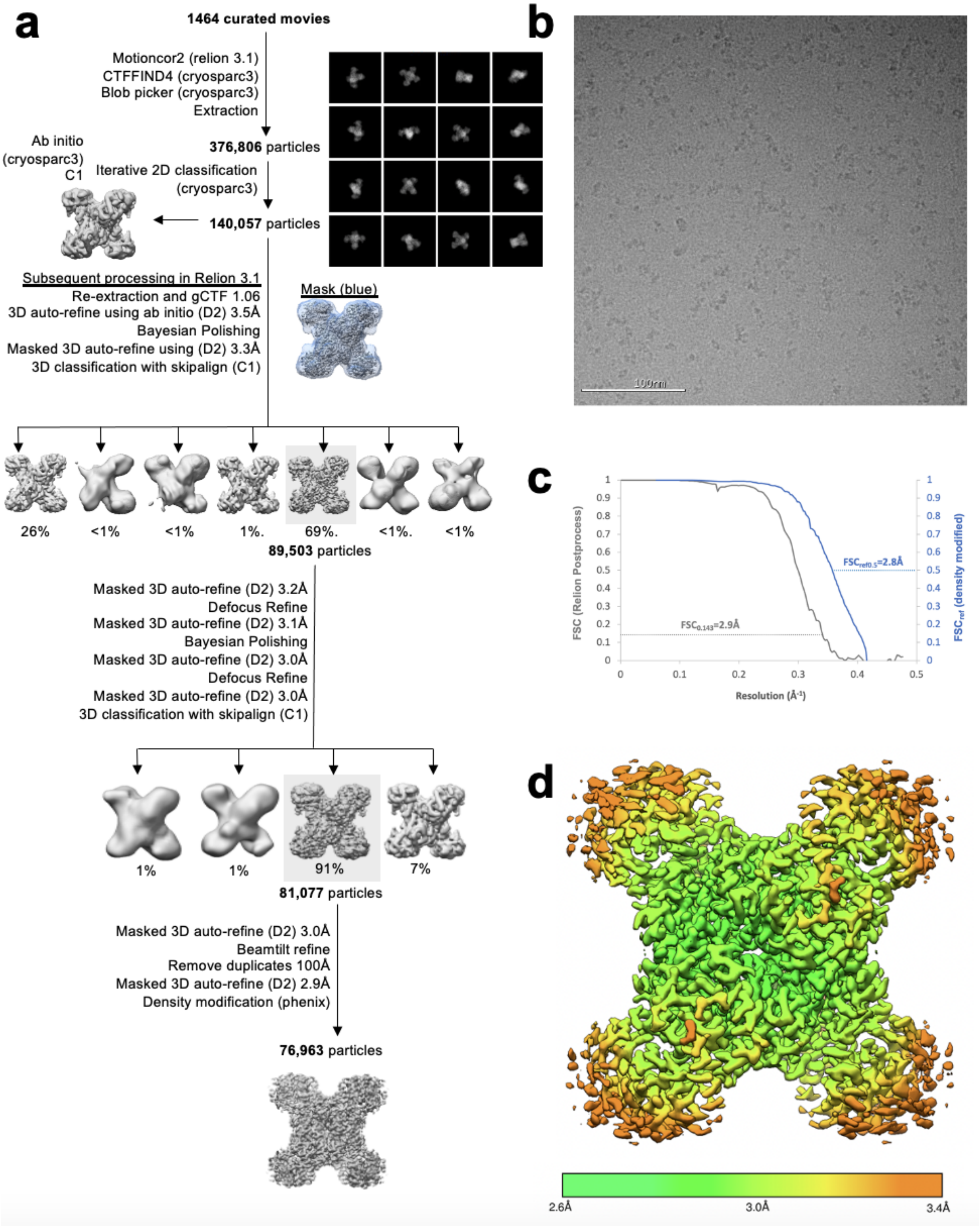


**Fig3. Supplemental 9. Image Processing of Substrate-bound Ura8 Tetramers.** (a) Flowchart overview of the data processing strategy. (b) example micrograph from the dataset. (c) FSC curve from Relion postprocess and Phenix density modification. (d) local resolution estimation.

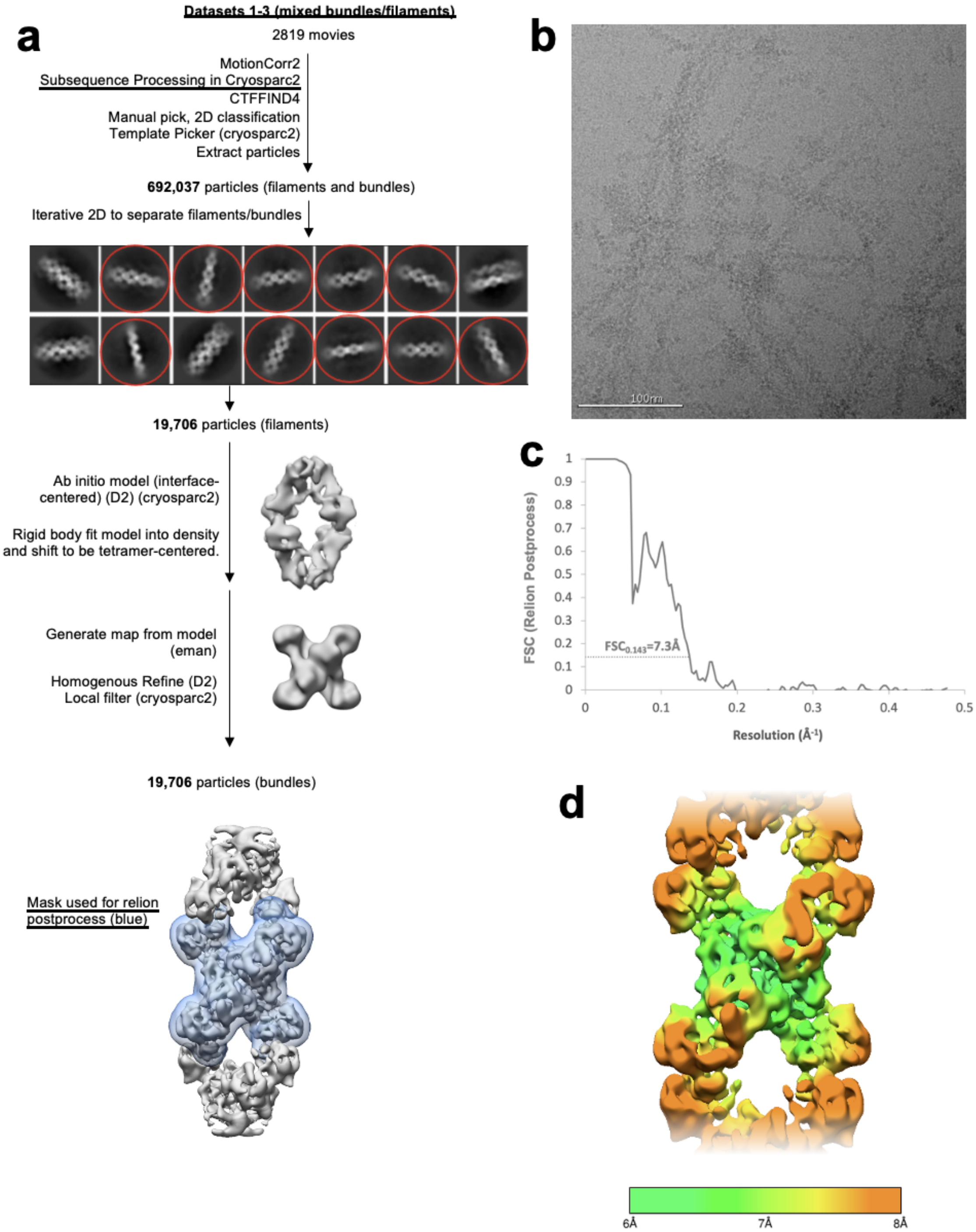


**Fig3. Supplemental 10. Image Processing of Substrate-bound Ura7 Filaments.** (a) Flowchart overview of the data processing strategy. (b) example micrograph from the dataset. (c) FSC curve from Relion postprocess. (d) local resolution estimation.

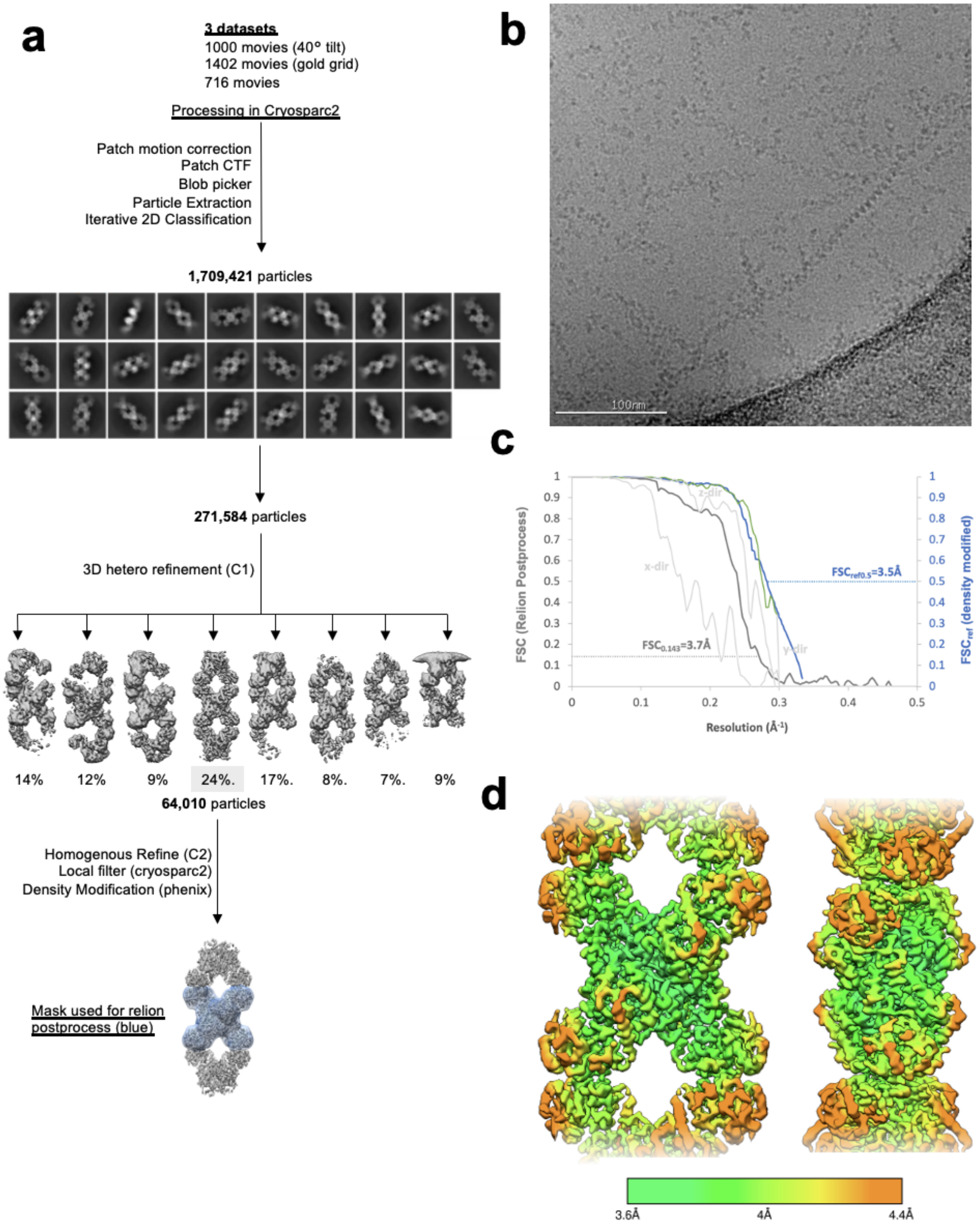


**Fig3. Supplemental 11. Image Processing of Product-bound Ura7 Filaments.** (a) Flowchart overview of the data processing strategy. (b) example micrograph from the dataset. (c) FSC curve from Relion postprocess and Phenix density modification. Light grey is 3D FSC for X/Y/Z directions. (d) local resolution estimation.

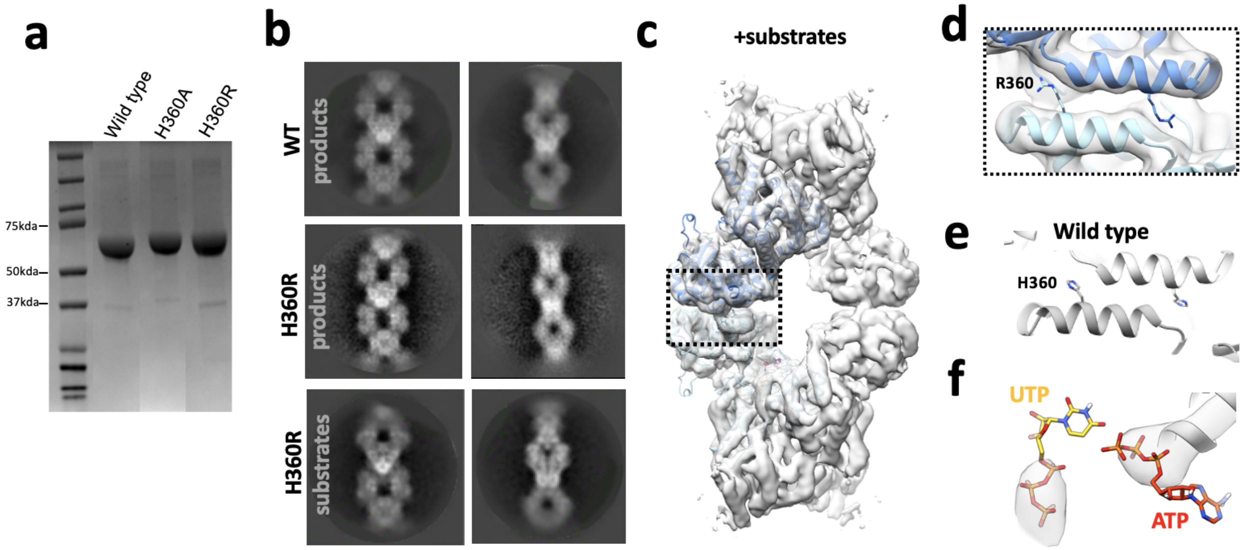


**Fig4. Supplemental 1. Ura7 H360R structural validation.** (a) SDS-PAGE gel of wild type and mutants purified in tandem. (b) representative 2D class averages of product-bound Ura7 wild type, and product or substrate-bound Ura7 H360R. (c) 3D reconstruction of substrate-bound Ura7 H360R with two monomers placed by rigid body fit of domains. (d) Zoom in from box in panel C of Ura7 H360R assembly interface with arginines modelled at residue 360. (e) wild type Ura7 interface. (f) nucleotide density from reconstruction in panel c.

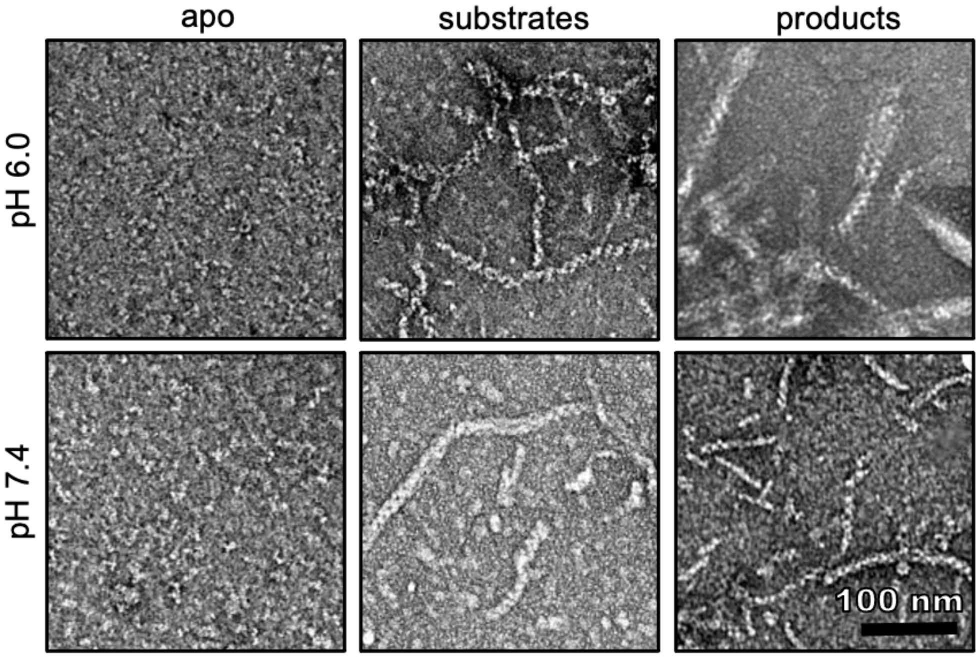


**Fig4. Supplemental 2. Ura7 H360R Ligand dependent Assembly.** Negative stain EM of Ura7 H360R assembly at low and high pH with addition of substrates (2mM UTP/2mM ATP) or product (2mM CTP).

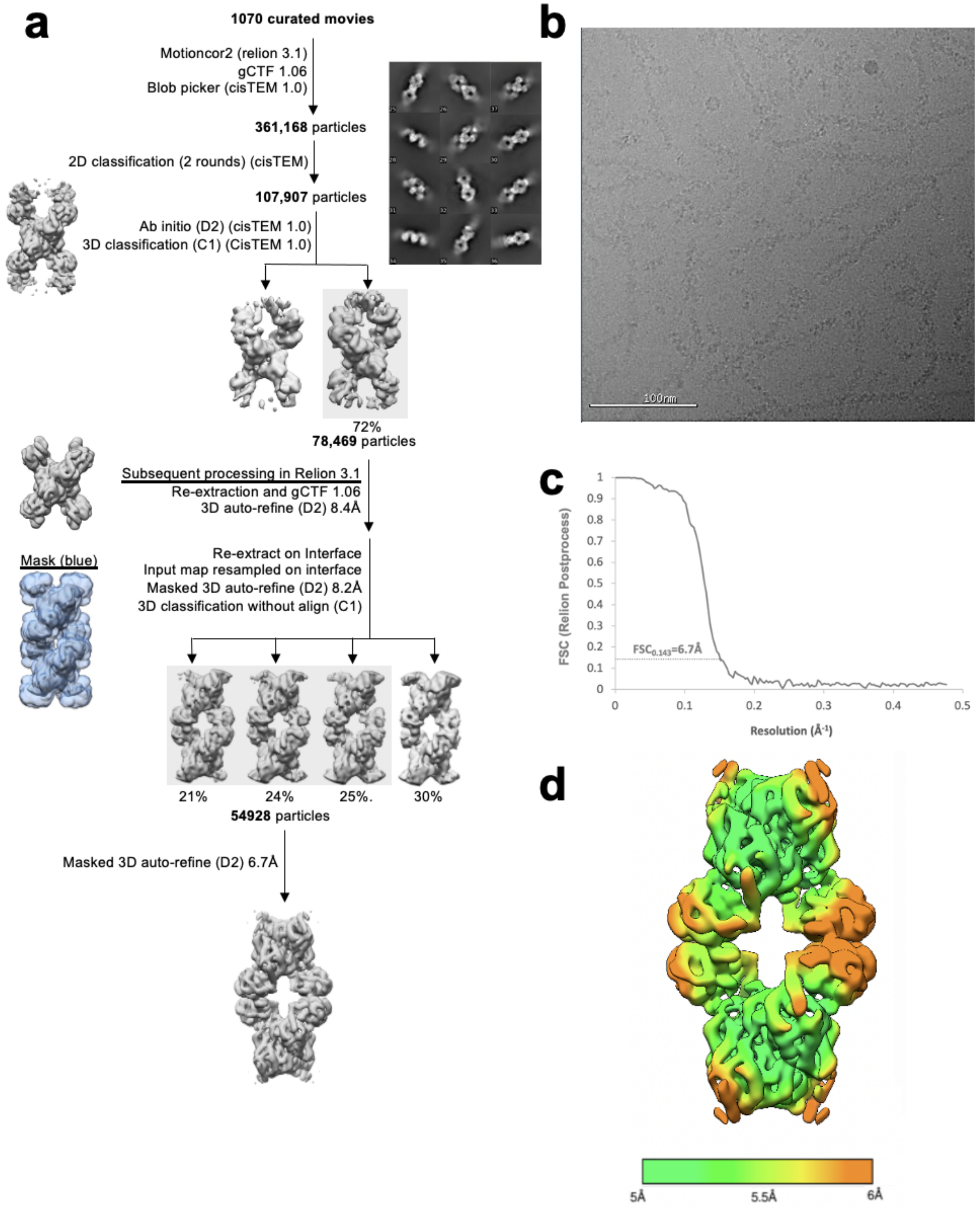


**Fig6. Supplemental 6. Image Processing of Substrate-bound Ura7-H360R at pH 7.5.** (a) Flowchart overview of the data processing strategy. (b) example micrograph from the dataset on continuous carbon. (c) FSC curve from Relion postprocess. (d) local resolution estimation.

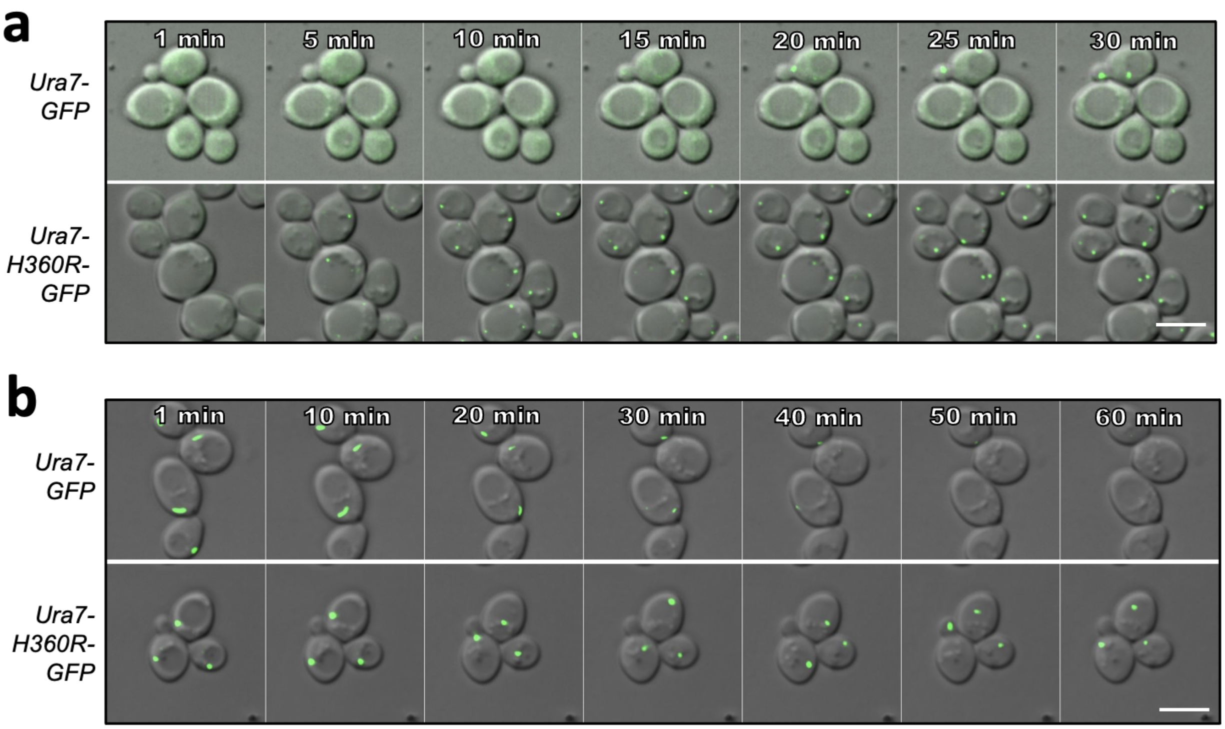


**Figure 5 supplemental 1. Ura7 assembly and disassembly kinetics (a)** Yeast expressing GFP-tagged Ura7 upon transfer from log-phase growth conditions to minimal starvation media (at time 0). Scale bar is 5 microns. **(b)** Yeast expressing GFP-tagged Ura7 after 4 hours starvation upon transfer to nutrient rich media. Scale bar is 5 microns.

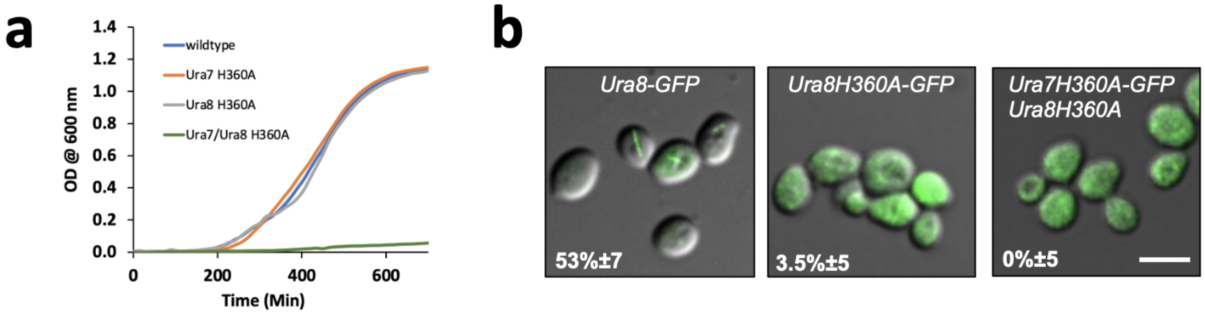


**Figure 5 supplemental 2. URA8 Non-assembly mutant.** (a) liquid culture growth curve for wild type and mutant Ura7 and Ura8. (b) GFP-tagged Ura8 wild type and mutant after 4 hours of starvation. Scale bar is 5 microns.

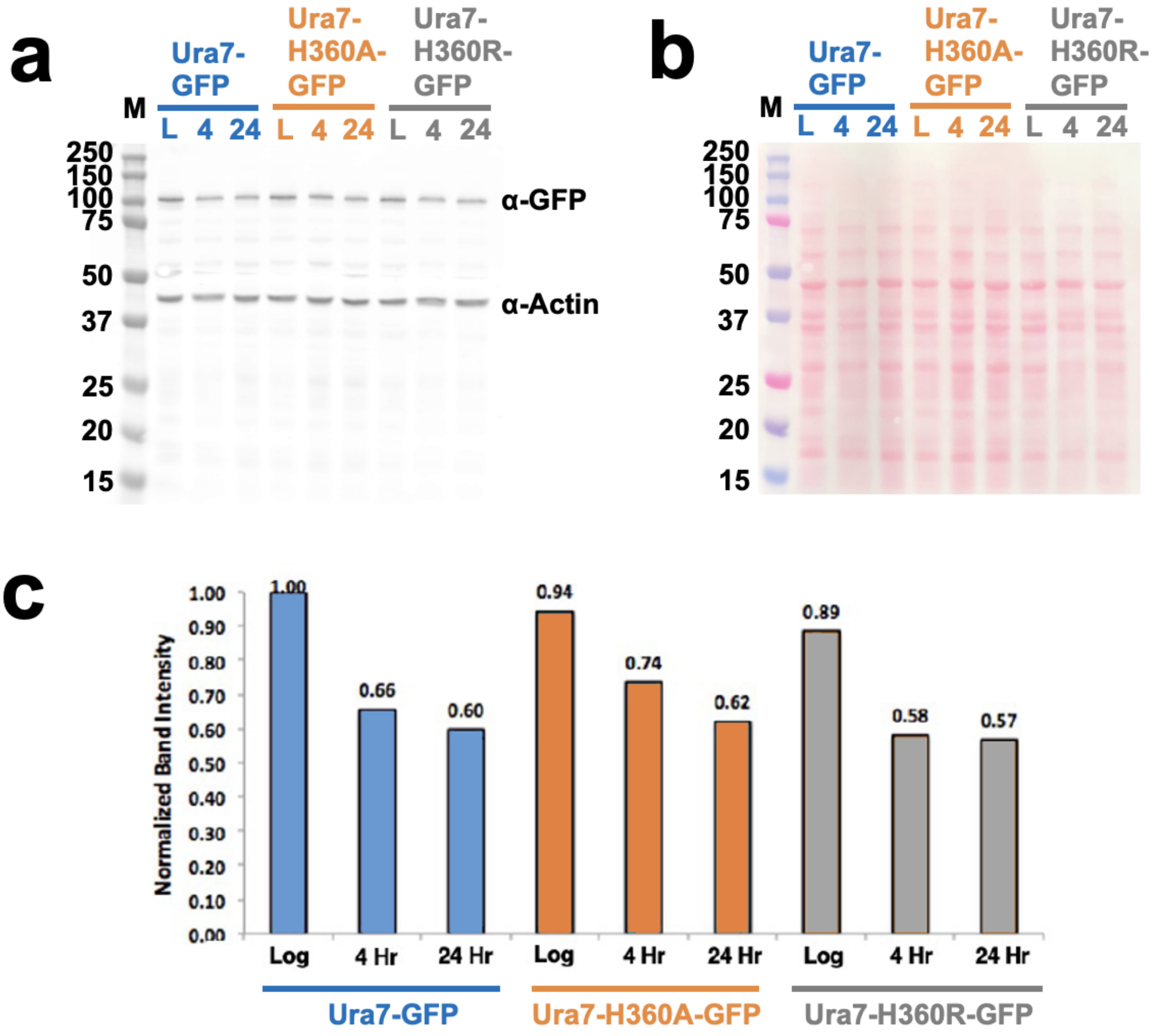


**Figure 5 supplemental 3. Ura7 Quantification from Yeast Cell Lysates.** (a) GFP-Ura7 expressed in yeast and quantified by Western blot. Wells were normalized by OD600 of cells, and Ura7 levels were assessed by anti-GFP antibody with anti-actin control. Ura7 was assessed in log phase ("L"), and after 4 or 24 hours of starvation. Ladder, left, is in kilo-dalton. (b) Ponceau stain of western blot from panel a. (c) quantification of band intensity from panel A, relative to the most intense band, are displayed above each bar.

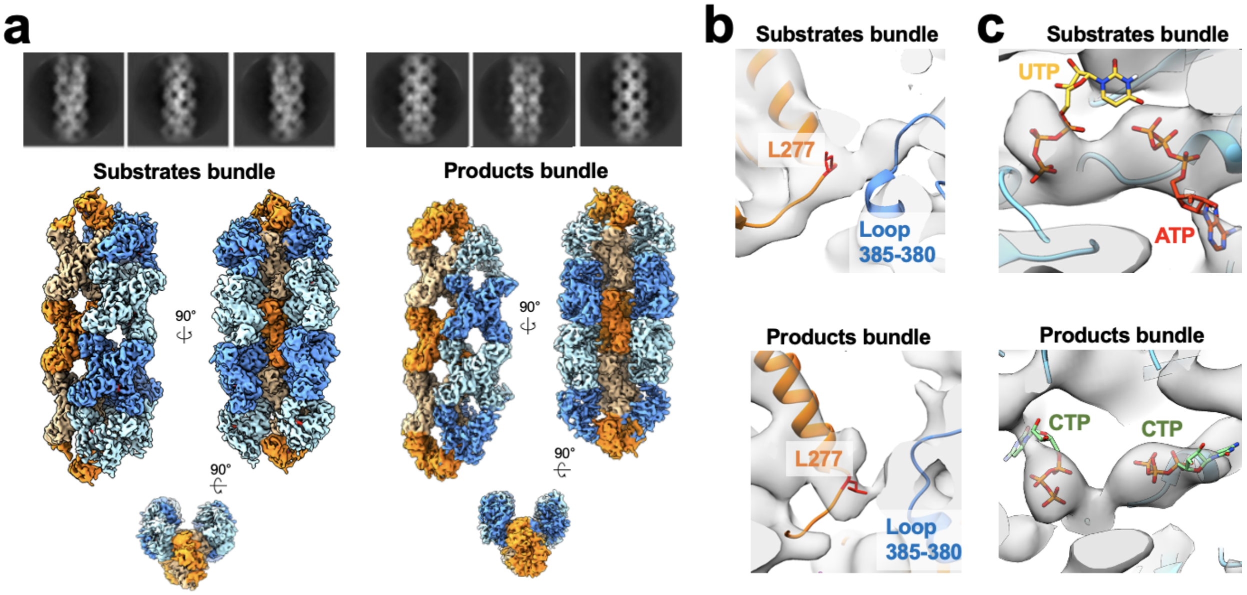


**Fig6. Supplemental 1. Ura7 Bundle architecture. (a)** Ura7 substrate- (2mM UTP/2mM ATP) and product- (2mM CTP) bound cryo-EM 2D averages and 3D reconstructions. **(b)** Lateral contact mediated by yeast linker region (orange) with adjacent strand. Residue 277 is highlighted in red (see text). **(c)** Nucleotides in the active site with corresponding density.

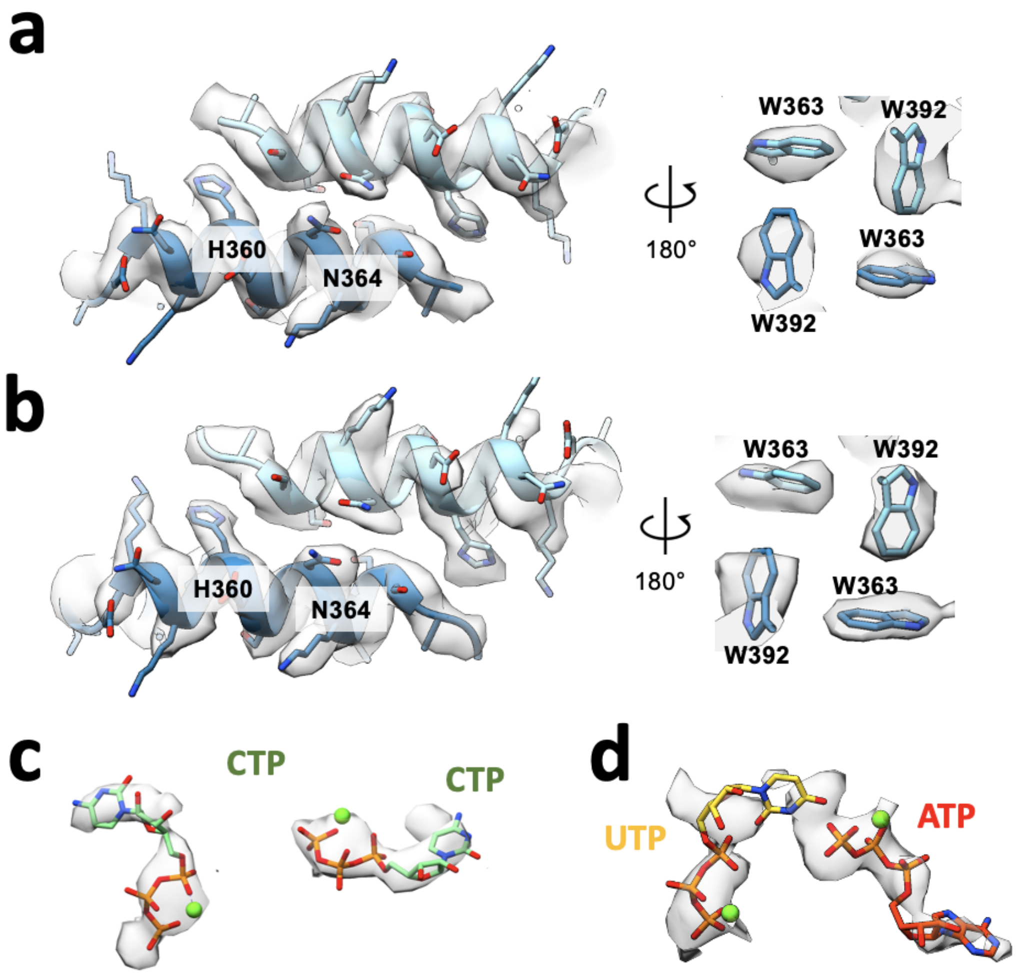


**Fig6. Supplemental 2. Filament assembly interface and nucleotides for Ura8 in bundles (a)** Assembly interface and corresponding density for an individual product-bound Ura8 strand while in a bundle. Key residues are indicated. **(b)** Assembly interface as in A, except for substrate-bound Ura8 bundle. **(c-d)** Nucleotides and corresponding density for substrate (c; 2mM UTP/2mM ATP) and product-bound (d; 2mM CTP) bundle reconstructions.

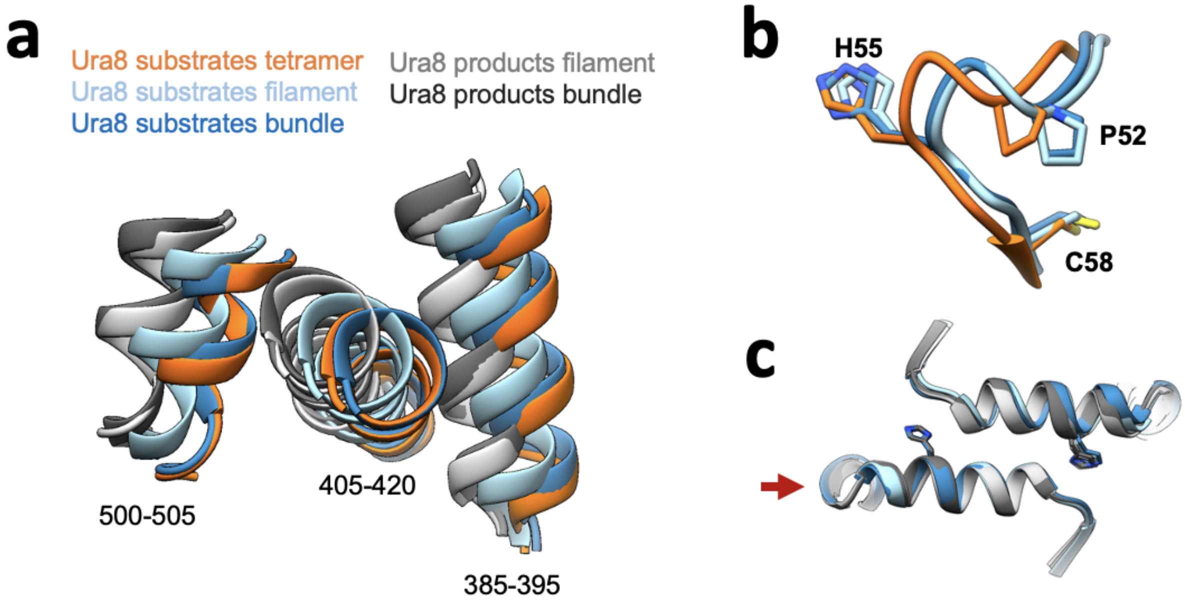


**Fig6. Supplemental 3. Comparison of filaments from individual filaments versus strands within bundles.** **(a)** Glutaminase domain rotation of helices shown in figure 4 (numbered here). **(b)** Ammonia channel key residues for closure for substrate-bound Ura8 bundle relative to tetramer and filament state. **(c)** Filament assembly interface for Ura8 either in bundles or individual filaments. Red arrow shows which side of the interface models were aligned on.

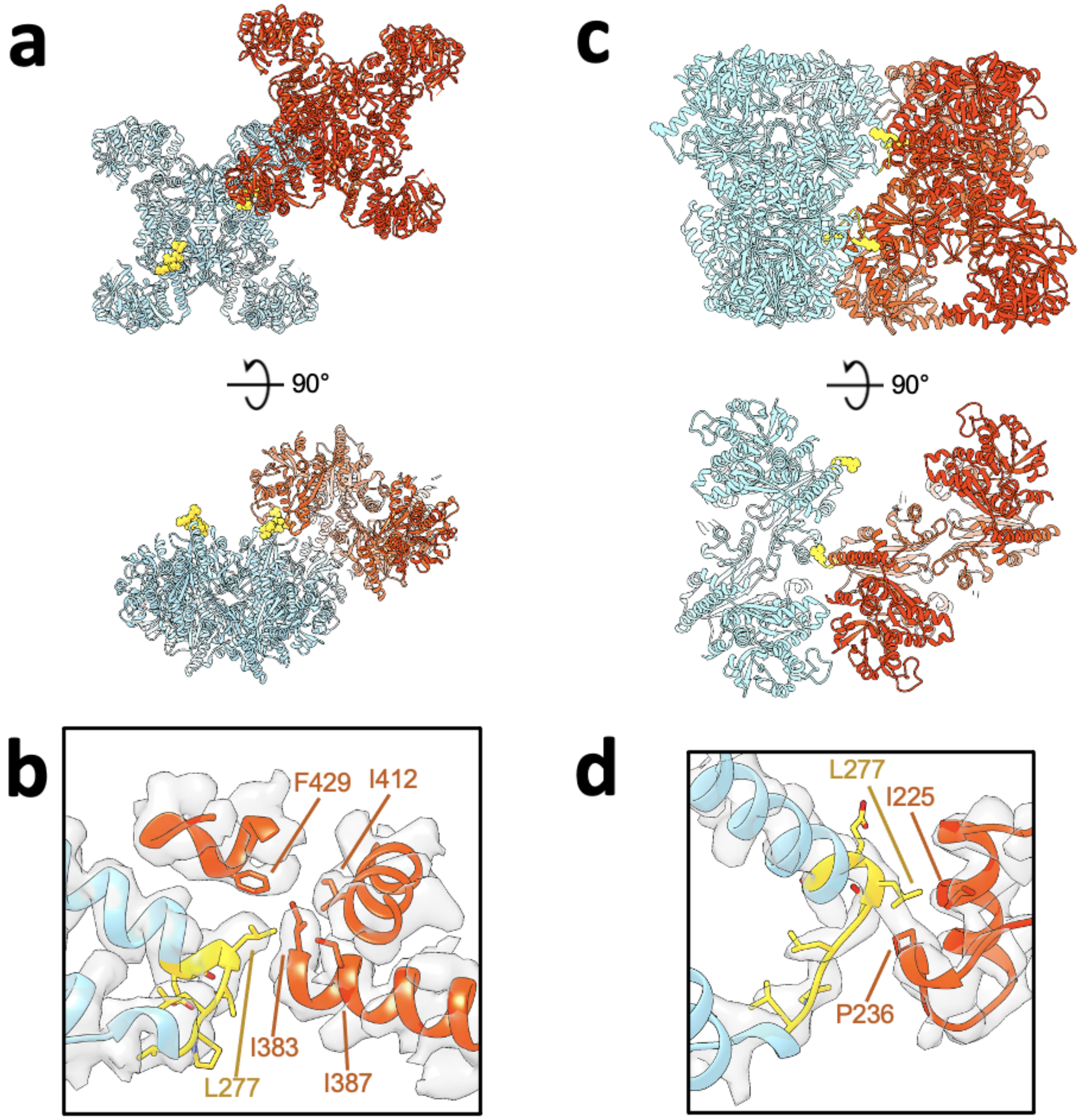


**Fig6. Supplemental 4. Yeast Linker Involvement in Lateral Bundle Contacts (a)** Substrate-bound Ura8 bundle, shown are a pair of tetramers interacting laterally. Colored in yellow is the yeast linker insert (residues 273-279). **(b)** zoom in of yeast linker insert contact with corresponding cryo-EM density. Yeast Insert residues putative contacts on the opposing strand are displayed. **(c-d)** same as a-b but for a pair of tetramers in the product-bound Ura8 bundle.

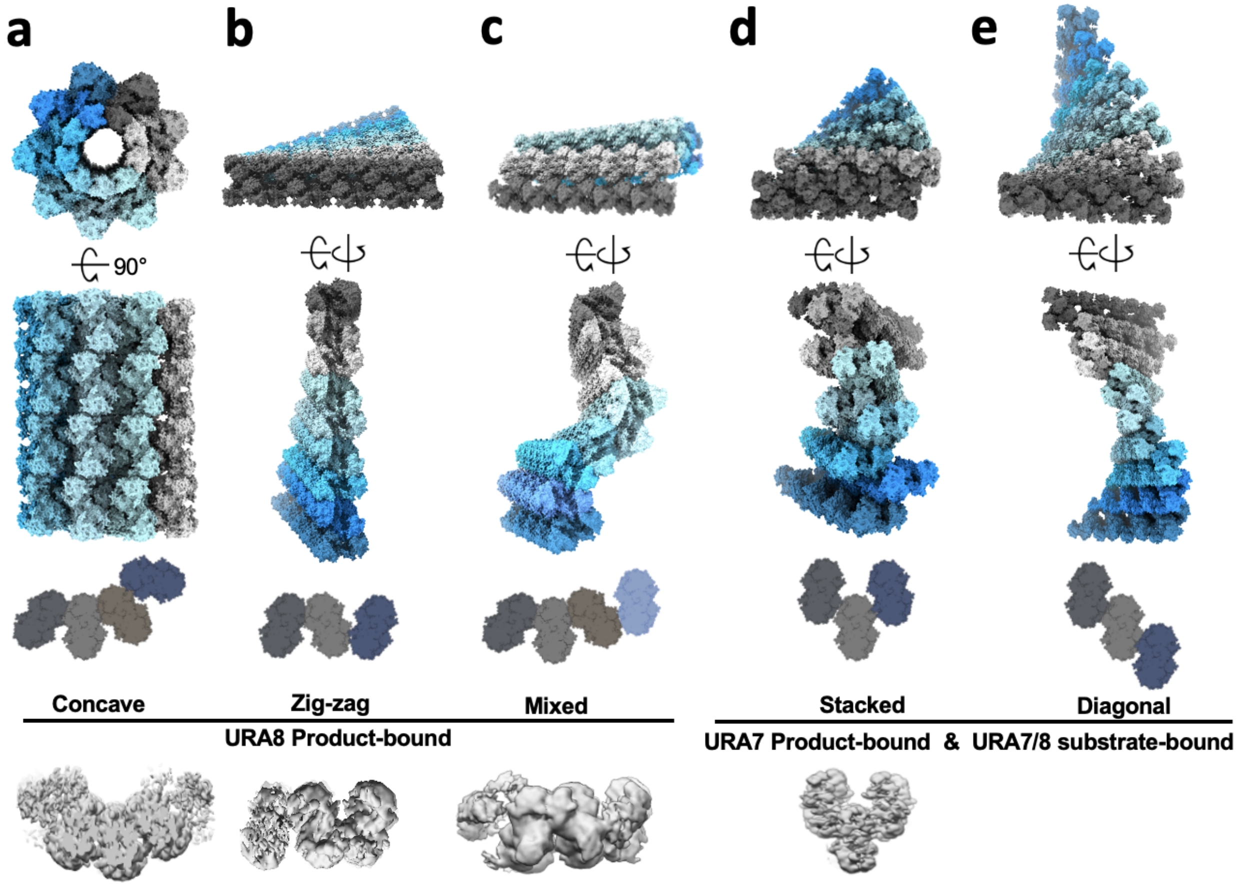


**Fig6. Supplemental 5. Possible higher-order arrangements of yeast CTPS bundles.** Models of possible bundle configurations generated in silico by propagating contacts. Bottom shows end-on views of 3D classification maps done in C1 which were used as templates to generate the propagated models above. (**a-c**) Configurations for product-bound Ura8 which has strands in register. **(d-e)** Configurations for staggered strand architecture adopted by Ura8 substrate-bound and Ura7 in both ligand states. No 3D map was obtained for a diagonal configuration but our modelling predicts that it should be possible without clashing.

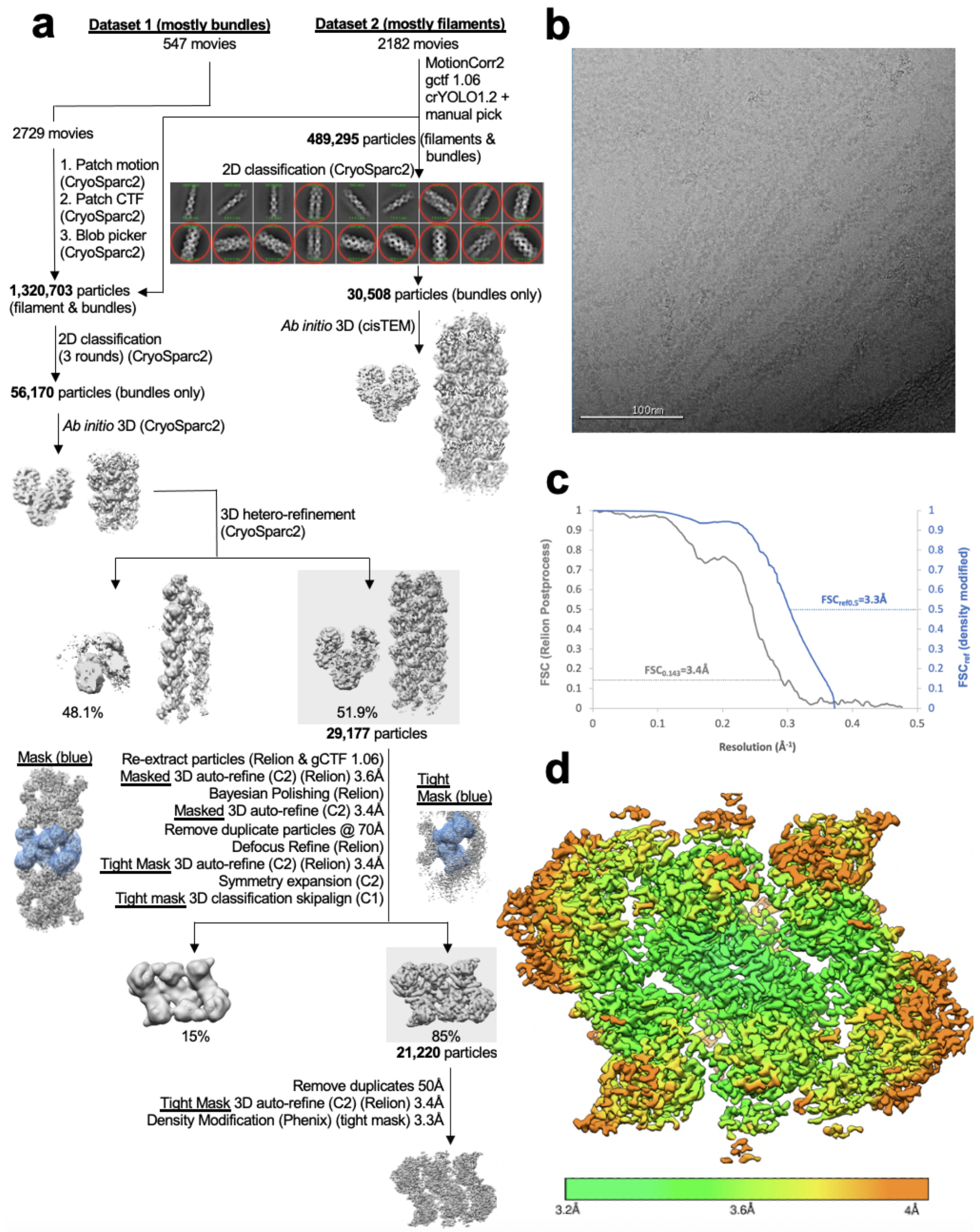


**Fig6. Supplemental 6. Image Processing of Substrate-bound Ura8 Bundles.** (a) Flowchart overview of the data processing strategy. (b) example micrograph from the dataset. (c) FSC curve from Relion postprocess and Phenix density modification. (d) local resolution estimation.

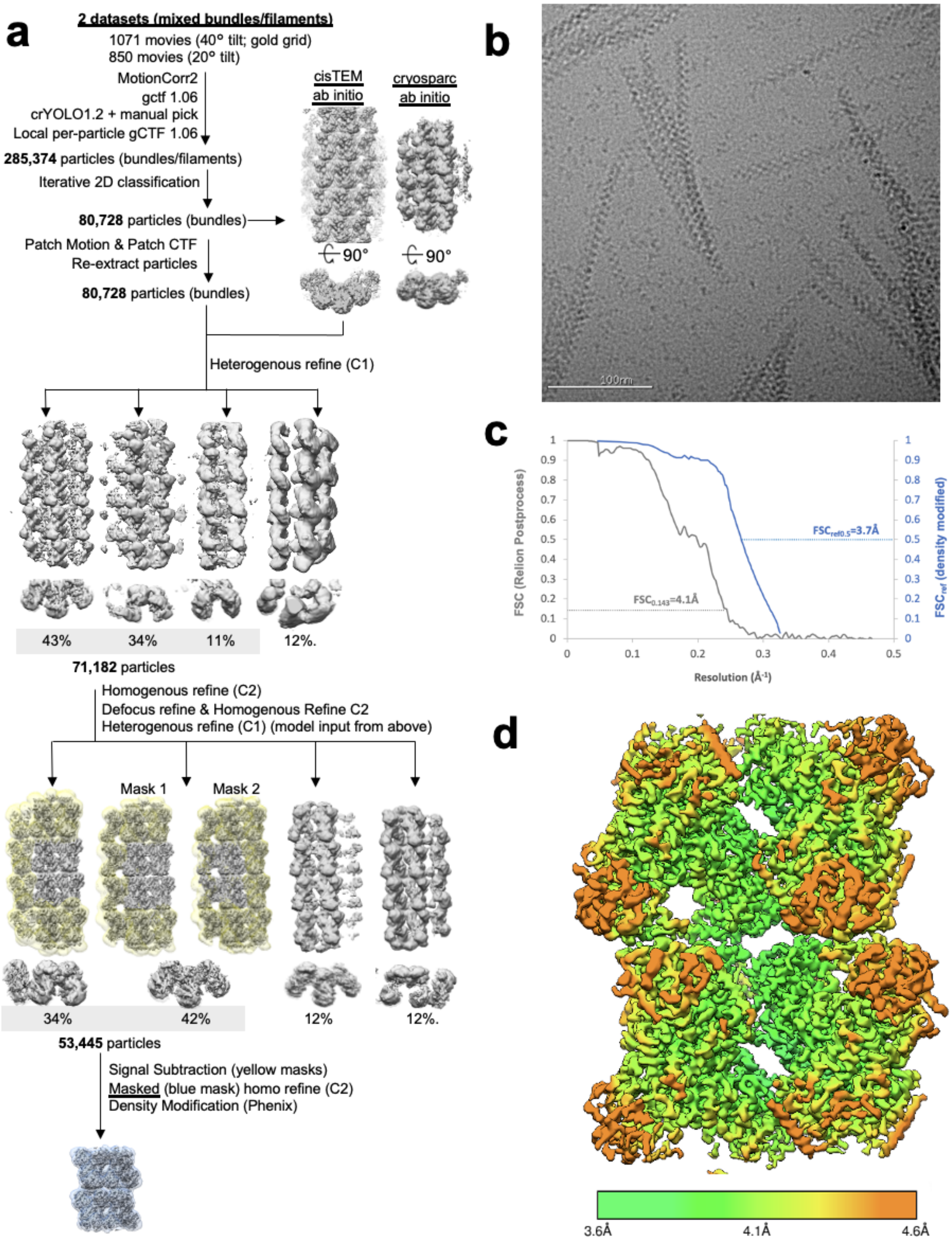


**Fig6. Supplemental 7. Image Processing of Product-bound Ura8 Bundles.** (a) Flowchart overview of the data processing strategy. (b) example micrograph from the dataset. (c) FSC curve from Relion postprocess and Phenix density modification. (d) local resolution estimation.

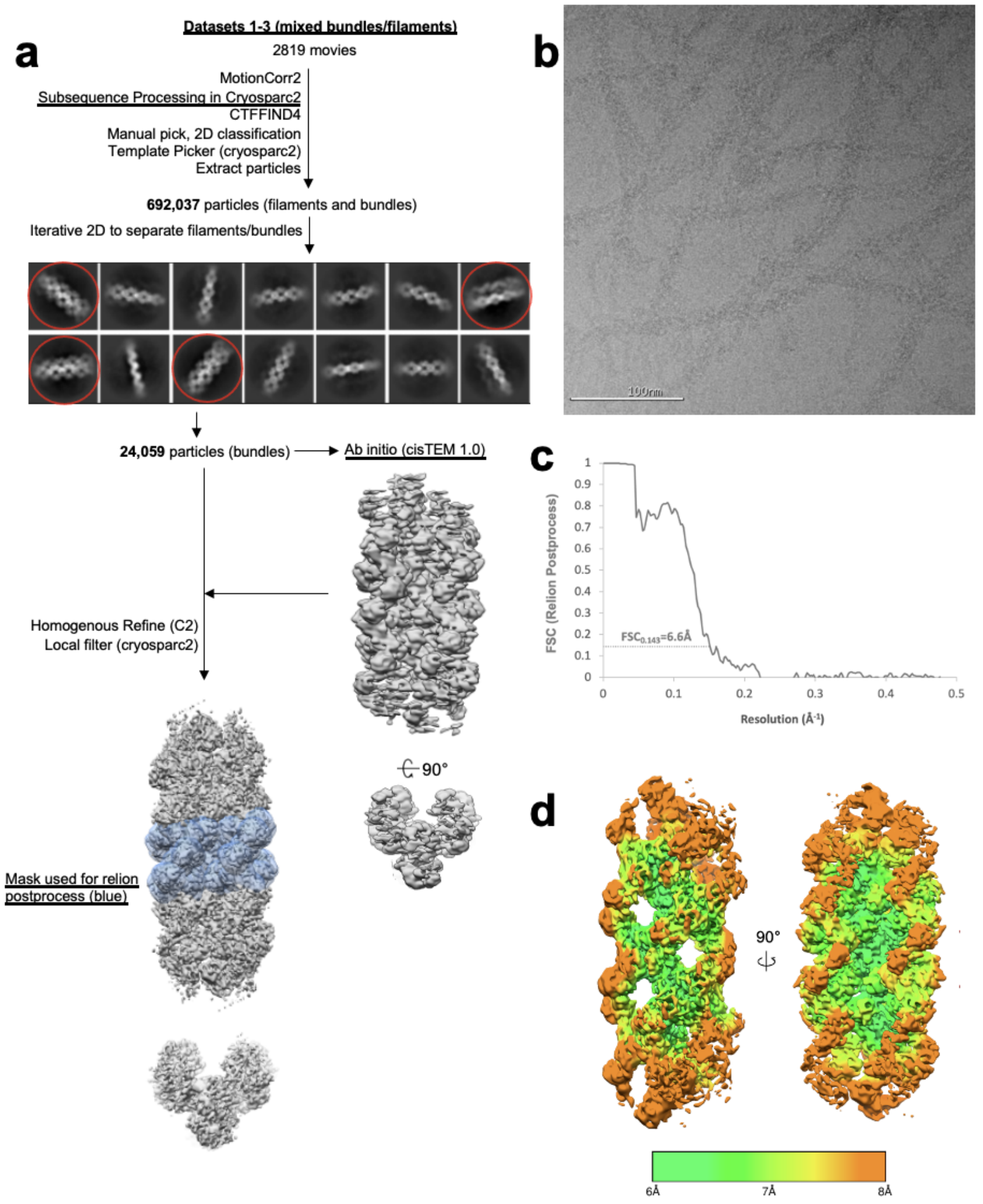


**Fig6. Supplemental 8. Image Processing of Substrate-bound Ura7 Bundles.** (a) Flowchart overview of the data processing strategy. (b) example micrograph from the dataset. (c) FSC curve from Relion postprocess. (d) local resolution estimation.

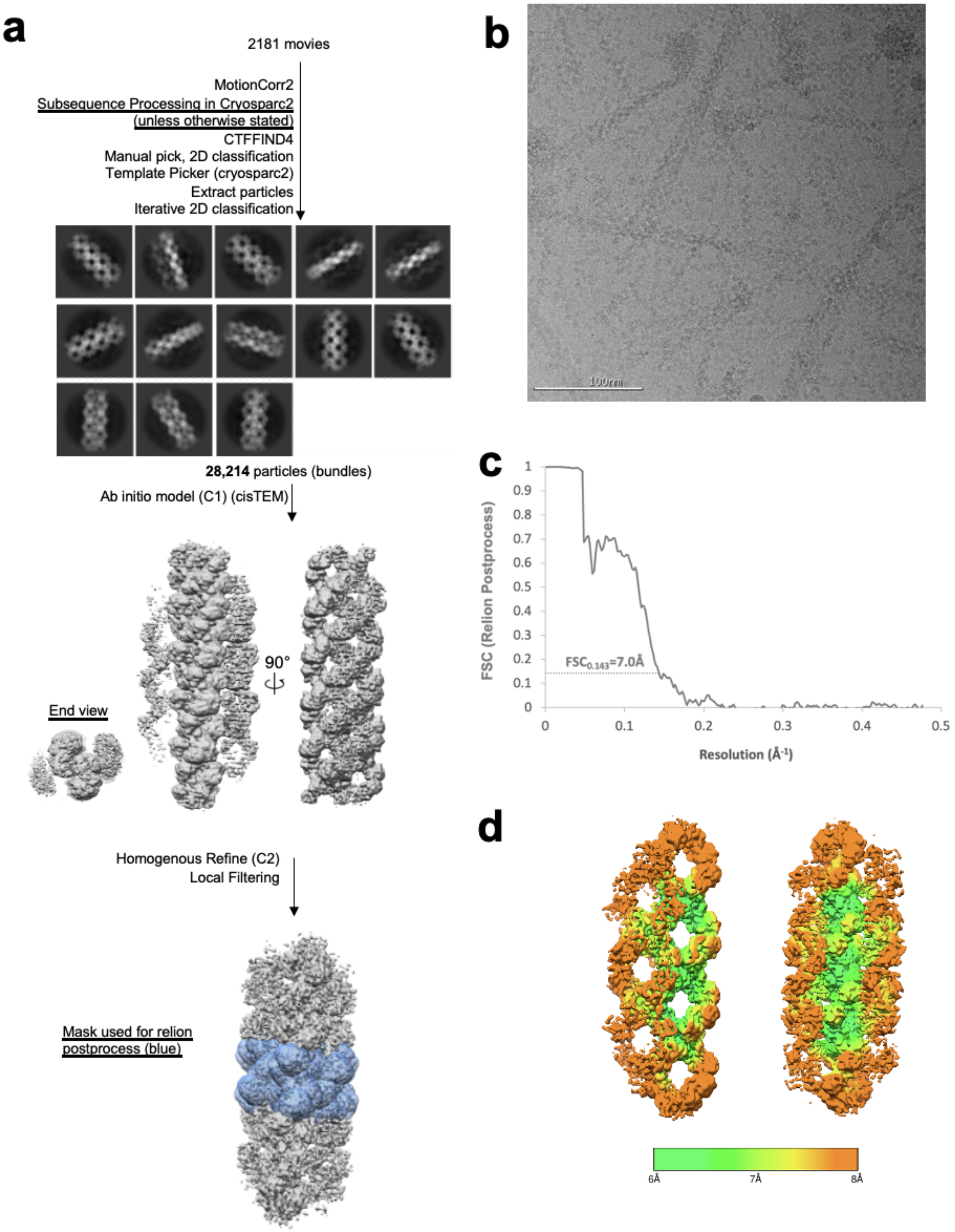


**Fig6. Supplemental 9. Image Processing of Product-bound Ura7 Bundles.** (a) Flowchart overview of the data processing strategy. (b) example micrograph from the dataset. (c) FSC curve from Relion postprocess. (d) local resolution estimation.

### Table 1. Cryo-EM Data Collection and Refinement Statistics

| | Ura8 substrates filament | Ura8 Substrates Bundle | Ura8 substrates tetramer | Ura8 products filament | Ura8 products bundle | Ura7 substrates filament | Ura7 substrates bundle | Ura7 products filament | Ura7 products bundle | Ura7-H360R substrates filament |
|---|---|---|---|---|---|---|---|---|---|---|
| Number of micrographs | 2729 | 2729 | 1464 | 4979 | 1921 | 2819 | 2819 | 3118 | 2181 | 1070 |
| Nominal magnification | 130,000X | | | | | | | | | |
| Voltage | 300 kV | | | | | | | | | |
| Electron Fluence | $90e^-/Å^2$ | | | | | | | | | |
| Pixel size | 1.05Å | | | | | | | | | |
| Defocus range | -0.4 to -1.9μm | -0.4 to -1.9μm | -1.2 to -5.5μm | -0.4 to -9.0μm | -0.4 to -7.0μm | -0.8 to -2.2μm | -0.8 to -2.2μm | -0.4 to -7.0μm | -1.0 to -2.2μm | -0.9 to -3.5μm |
| EMDB ID | EMD-24512 | EMD-24581 | EMD-24497 | EMD-24516 | EMD-24579 | EMD-24566 | EMD-24575 | EMD-24560 | EMD-24576 | EMD-24578 |
| PDB code | 7RL0 | 7RNR | 7RKH | 7RL5 | 7RNL | 7RMF | 7RMK | 7RMC | 7RMO | 7RMV |
| Map resolution 0.143 FSC | 2.9Å | 3.4Å | 2.9Å | 4.0Å | 4.1Å | 7.3Å | 6.6Å | 3.7Å | 7.0Å | 6.7Å |
| Density Modified Resolution 0.5Ref | 2.8Å | 3.3Å | 2.8Å | 3.8Å | 3.7Å | n/a | n/a | 3.5Å | n/a | n/a |
| Symmetry Imposed | D2 | C2 | D2 | D2 | C2 | D2 | C2 | D2 | C2 | D2 |
| Number of particles | 40,474 | 21,220 | 76,963 | 181,136 | 53,445 | 19,706 | 24,059 | 64,010 | 28,214 | 54,928 |
| Map Sharpening B Factor | $-32Å^2$ | $-33Å^2$ | $-74Å^2$ | $-142Å^2$ | $-67Å^2$ | $-235Å^2$ | $-140Å^2$ | $-51Å^2$ | $-186Å^2$ | $-40Å^2$ |
| **Model composition** | | | | | | | | | | |
| ligands | ATP/UTP | ATP/UTP | ATP/UTP | CTP | CTP | ATP/UTP | ATP/UTP | CTP | CTP | ATP/UTP |
| **R.M.S. deviations** | | | | | | | | | | |
| Bond lengths (Å) | 0.64 | 0.63 | 0.63 | 0.66 | 0.7 | 0.69 | 0.69 | 0.62 | 0.63 | 0.71 |
| Bond angles (°) | 1.09 | 1.04 | 1.04 | 1.09 | 1.13 | 1.06 | 1.06 | 1.03 | 1.03 | 1.07 |
| **Validation** | | | | | | | | | | |
| Molprobity Score | 0.98 | 1.35 | 1.09 | 1.59 | 1.66 | 1.56 | 2.06 | 1.40 | 1.73 | 2.30 |
| Clash Score | 0.94 | 4.74 | 0.9 | 3.2 | 3.48 | 2.31 | 10.33 | 1.51 | 4.62 | 15.84 |
| Poor Rotamers (%) | 1.21 | 0.61 | 1.64 | 2.05 | 1.43 | 1.88 | 1.88 | 1.65 | 1.66 | 2.09 |
| Ramachandran Plot | | | | | | | | | | |
| Favored (%) | 97 | 98 | 97 | 96 | 94 | 95 | 95 | 95 | 95 | 95 |
| Allowed (%) | 2 | 2 | 3 | 3 | 6 | 4 | 4 | 5 | 4 | 5 |
| Outliers (%) | 0 | 0 | 0 | 0 | 0 | 0 | 0 | 0 | 0 | 0 |

**Table 2. CTPS Polymer Architecture Characteristics**

| | Ura8 prods filament | Ura8 subs filament | Ura8 subs tetramer | Ura8 prods bundle | Ura8 subs bundle | Ura7 prods filament | Ura7 Subs filament | Ura7 prods bundle | Ura7 subs bundle |
|---|---|---|---|---|---|---|---|---|---|
| **Domain rotation relative to product-bound filament state** | **n/a** | **4°** | **7.4°** | **1.7°** | **6.9°** | **n/a** | **7.7°** | **1.4°** | **6.2°** |
| **Filament twist** | **5°** | **17.1°** | **n/a** | **3.9°** | **15.9°** | **7.4°** | **18.2°** | **7.3°** | **17.3°** |
| **Filament rise** | **102.6Å** | **102.3Å** | **n/a** | **103Å** | **101.8Å** | **102.7Å** | **103.4Å** | **102.8Å** | **104.5Å** |

**Table 3. Kinetic activity parameters for yeast CTPS and mutants.**

| | Wild type | H360A | H360R |
|---|---|---|---|
| **Max** | 2.68 | 3.60 | 1.18 |
| **Min** | 0.00 | 0.00 | 0.00 |
| **S50** | 161.07 | 160.03 | 229.72 |
| **Hill** | 1.53 | 1.45 | 0.80 |

**Table 4. CTPS Bundle Buried Surface Area and Putative Contact Residues.**

| **Ura8 Product-bound Bundle Contacts** | | | | | | |
|---|---|---|---|---|---|---|
| **BSA** | **domains** | **Monomer 0** | **Monomer 1** | **Monomer 2** | **Monomer 3** | **Monomer 4** |
| 93Å$^2$ | Glu-Glu | 465,468,515 | 381, 384, 429,430,432,433 | | | |
| 173Å$^2$ | Glu-Glu | 471,472,476,477†,480-483 | 348,350†,352, 353† | | | |
| 113Å$^2$ | Glu-Glu | 566,567*,570, 571 | 354-356,358 | | | |
| 240Å$^2$ | Link-AL | 275,276,277*, 278,280†,281† | | 222†,225,226*,230,236,237† | | |
| 48Å$^2$ | Link-AL | 272,274 | | | 168,169,171,172 | |
| **Ura8 Substrate-bound Bundle Contacts** | | | | | | |
| 471Å$^2$ | Link-Glu | 275,276,277*, 278-282,284,285,288 | | | 348,350,380-383,384*,387, 412,416,423,425,426,429,430,432,433,435 | |
| 98Å$^2$ | AL-Glu | 136-139 | | | 353,354 | |
| 185Å$^2$ | Glu-AL | 471,472,477,481,506,507,526,527 | | 168-172,174,272,274 | | |
| 152Å$^2$ | Glu-AL | 563,567,570,571† | | 100†, 105-107,109,119,123, 126,127,130 | | |
| 88Å$^2$ | Glu-AL | 465,468,471,476,483,484 | 233,236-238,263 | | | |
| 151Å$^2$ | AL-Glu | 168-172,174 | | | | 469,471,472,477 |
| 167Å$^2$ | AL-Glu | 100,105-107, 123†,126*, 130† | | | | 561,563,564†, 567†,571 |

* predicted hydrogen bond. † predicted salt bridge

**Table 5. Yeast Strains Used**

| Number | Background | Genotype |
|---|---|---|
| yJMK001 | W303 | URA7-GFP::KanMx |
| yJMK002 | W303 | URA7H360A |
| yJMK003 | W303 | URA7H360A-GFP::KanMx |
| yJMK004 | W303 | URA7H360R |
| yJMK005 | W303 | URA7H360R-GFP::KanMx |
| yJMK006 | W303 | URA8-GFP::KanMx |
| yJMK007 | W303 | URA8-mCherry::KanMx |
| yJMK008 | W303 | URA8H360A |
| yJMK009 | W303 | URA8H360A-GFP::KanMx |
